# Construction of a GnRH mRNA Immunocastration Vaccine and Evaluation of Its Immunogenicity and Safety in Mice and Cats

**DOI:** 10.64898/2026.03.25.714088

**Authors:** Yue Chen, Chao Dong, Wenqian Yan, Yujing Liu, Jixiang Sun, Mingshuo Ji, Jiahan Gang, Jiaqi Nie, Xinhao Zhang, He Huang, Yongfei Zhou

**Author notes:** These authors contributed equally to this work (Yue Chen, Chao Dong, Wenqian Yan, Yujing Liu). **Corresponding author:** Yongfei Zhou, **Email addresses:**.

## Abstract

Immunocastration has emerged as an alternative to surgical and chemical castration for managing reproductive function in animals, yet the development of safe and effective vaccines remains challenging. This study aimed to develop a gonadotropin-releasing hormone (GnRH)-based messenger RNA (mRNA) vaccine and systematically evaluate its immunogenicity, reproductive suppression efficacy, long-term durability, and biosafety in mice and cats. GnRH epitopes were fused to three carrier proteins, Fc, Foldon, and lumazine synthase nanoparticles (pLS) via a flexible linker. After identifying pLS as the optimal scaffold, three mRNA vaccine candidates (GnRH-3, GnRH-4, and GnRH-5) were generated with one, five, or ten tandem GnRH repeats, encapsulated in lipid nanoparticles (LNPs), and assessed in rodent and feline models. Immunogenicity was determined by enzyme-linked immunosorbent assay, gonadal histopathology, hormone measurements, transcriptomic analysis, and mating trials. Among the fusion partners, the pLS-based vaccine (GnRH-3) induced the strongest antibody responses and most pronounced reproductive suppression. Further optimization showed that GnRH-4, containing five tandem GnRH repeats, elicited the highest antibody titers, induced severe gonadal atrophy, and reduced litter size by 93.8% in mice. Transcriptomic analysis revealed that differentially expressed genes in males were enriched in spermatogenesis and motility pathways, whereas those in females were associated with RNA splicing and immune responses. In cats, the optimal regimen was a twoLdose schedule with 50 μg per dose and a 21Lday interval, which induced robust antibody responses lasting at least 12 Lmonths and sustained reproductive suppression. HighLdose (500 μg) administration showed no clinical toxicity or histopathological abnormalities, confirming favorable biosafety. This study successfully developed a pLSLbased GnRH mRNA vaccine (GnRH-4) with five tandem GnRH epitopes that demonstrates strong immunogenicity, longLlasting contraceptive effects, and excellent safety in both rodent and feline models, supporting its potential for clinical application in immunocastration.

## 1. Introduction

Castration is a long-standing practice in animal husbandry, offering advantages such as reducing aggressive behavior in male animals, improving meat quality, enhancing feed conversion efficiency, and managing sex hormone-dependent diseases [1]. It is also used to control wildlife populations and improve companion animal behavior. Traditional castration methods primarily include surgical castration and chemical castration. Surgical castration achieves permanent sterility through the surgical removal of the testes or ovaries but is associated with biological stress, wound infection, and postoperative acute pain [2, 3]. Chemical castration involves the administration of hormonal analogues such as GnRH agonists or antagonists to regulate fertility; although convenient, long-term administration may lead to drug resistance [4, 5]. In this context, immunocastration, which induces the production of specific antibodies to block reproductive hormone signaling pathways, thereby inhibiting gonadal development and reproductive function, offers advantages including simplicity, reversibility, and high safety [6, 7], and is considered an alternative to traditional castration.

Follicle-stimulating hormone (FSH) and luteinizing hormone (LH) are essential for normal sexual maturation and reproductive function in mammals. Upon entering the peripheral circulation, they act on the ovaries and testes to regulate folliculogenesis, ovulation, spermatogenesis, and steroidogenesis [8]. Gonadotropin-releasing hormone (GnRH), a decapeptide released by hypothalamic GnRH neurons, is a key regulator of mammalian reproductive system development. GnRH-based immunocastration vaccines target GnRH as the antigen; upon vaccination, the induced antibodies block the binding of GnRH to its receptors on pituitary gonadotrophs, thereby inhibiting the synthesis and secretion of LH and FSH, ultimately affecting reproductive function [9, 10]. To date, vaccination with GnRH-based vaccines has been evaluated for contraceptive efficacy in multiple animal species [11–13].

To confer immunogenicity against a self-protein, GnRH-based castration vaccines have been developed by coupling GnRH with various carrier proteins [14]. M. Takahashi et al. conjugated GnRH to bovine serum albumin (BSA) and vaccinated female rats, resulting in decreased LH levels and ovarian atrophy [15]. C. Carelli et al. linked GnRH to muramyl dipeptide via a lysine bridge, leading to reduced sperm production and decreased fertility in male mice. However, chemically conjugated GnRH vaccines, in which the target antigen GnRH is a small self-peptide with weak intrinsic immunogenicity, also face challenges such as high batch-to-batch variability and complex manufacturing processes [16]. C. T. Hsu et al. integrated a DNA fragment encoding an exotoxin receptor-binding domain fused with 12 copies of GnRH (PEIa-GnRH12) into an expression vector and expressed the recombinant protein in Escherichia coli; vaccinated female rabbits generated high-titer antibodies against GnRH [17]. Jinshu Xu et al. vaccinated female mice with a recombinant protein comprising three GnRH copies linked to T-cell epitopes of the measles virus protein (MVP) and similarly observed high-titer anti-GnRH antibodies. Building on this background, we developed an mRNA vaccine using Lumazine synthase nanoparticles as a carrier [18]. By designing an antigen sequence in which multiple tandem repeats of GnRH peptides are connected to the carrier via a GGGGS linker, the mRNA vaccine expresses proteins in vivo that self-assemble into virus-like particle (VLP) structures, greatly enhancing immunogenicity. This vaccine demonstrated favorable immunogenicity, castration efficacy, and safety in both mice and cats, providing a new strategy for the development of future immunocastration vaccines.

## 2. Materials and methods

### 2.1 Antigen design

To obtain GnRH mRNA vaccine candidates with high expression efficiency and strong immunogenicity, this study conducted a two-stage sequence screening and optimization process.

In the first stage, the amino acid sequence of the GnRH epitope (EHWSYGLRPG) was linked via a flexible linker (GGGGS)L to the Fc fragment (Fc) [19], the Foldon trimerization domain (Foldon) [20], and Lumazine synthase nanoparticles (pLS), respectively, generating three mRNA sequences with distinct fusion partners (designated GnRH-1, GnRH-2, and GnRH-3). A FLAG tag sequence (DYKDDDDK) was added to the C-terminus of each carrier sequence to facilitate protein expression analysis.

In the second stage, based on the optimal carrier identified from the first round of screening (GnRH-3), the GnRH epitope amino acid sequence along with the flexible linker (GGGGS)L was tandemly repeated 1, 5, or 10 times, respectively, and then fused to pLS (designated GnRH-3, GnRH-4, and GnRH-5, respectively), followed by codon optimization.

### 2.2 Plasmid construction and mRNA synthesis

The optimized coding sequences were cloned into a pUC57-derived mRNA expression plasmid under thel control of a T7 promoter. The expression cassette was flanked by optimized untranslated regions (UTRs): a 5′UTR and a 3′UTR derived from sequences known to enhance mRNA stability and translational efficiency, such as those from the α-globin gene. Additionally, a synthetic polyA tail of approximately 100 nucleotides was incorporated downstream of the 3′UTR to further enhance transcript stability. The plasmid construct was linearized downstream of the expression cassette and served as a template for in vitro transcription (IVT) using T7 RNA polymerase (Yeasen, Cat:10618ES90). The IVT reaction was supplemented with the co-transcriptional capping agent Cap1-GAG m7G(5’)ppp(5’)(2’0MeA)pG (Yeasen, Cat: 10677ES60) to ensure a high-fidelity 5′ cap structure, and N1-methylpseudouridine (Yeasen, Cat: 13860-38-3) was incorporated fully in place of uridine to reduce innate immune recognition and enhance translational efficiency. Following transcription, the mRNA was purified using lithium chloride precipitation (Thermo Fisher, Cat: AM9480) to remove enzymatic reagents and abortive transcripts, and its integrity, purity, and molecular size were confirmed by analytical agarose gel electrophoresis.

### 2.3 LNP formulation and characterization

The mRNA was encapsulated into LNPs utilizing a precision microfluidic-based assembly approach [21]. The optimized LNP formulation comprised four functional lipid components: the ionizable cationic lipid Heptadecan-9-yl 8-((2-hydroxyethyl) (6-oxo-6-(undecyloxy) hexyl) amino) octanoate) (HUO) (Sinopeg, China, Cat: 2089251-47-6), which facilitates mRNA complexation and endosomal disruption, 1,2-distearoyl-sn-glycero-3-phosphocholine (DSPC) (Sinopeg, Cat: 816-94-4) as a structural phospholipid, cholesterol (Sinopeg, Cat: 57-88-5) to enhance membrane stability and fluidity, and Methoxypoly (ethylene glycol) dimyristoyl glycerol (DMG-PEG 2000) (Sinopeg, Cat: 160743-62-4) to provide steric stabilization and reduce nonspecific interactions.These components were combined at an empirically determined molar ratio of 50:10:38.5:1.5 [22]. During the assembly process, an aqueous phase containing mRNA in citrate buffer (pH 4.0) and an organic phase comprising the lipid mixture in ethanol were simultaneously introduced into a microfluidic chip at a controlled volumetric ratio of 3:1(aqueous:organic), This process enabled rapid mixing and immediate nanoparticle formation within the microfluidic device (Fluidiclab, China, Model: NP-S2). The resulting mRNA-LNP suspension was subjected to ultrafiltration against Tris-HCl (pH 7.4) to remove residual ethanol, achieve buffer exchange, and ensure colloidal stability.The hydrodynamic diameter of the purified LNPs was determined by dynamic light scattering using a Zetasizer Nano ZS instrument (Brookhaven). Morphological attributes and structural integrity of the LNPs were visualized by transmission electron microscopy (TEM) (Hitachi, Japan).

### 2.4 Experimental Design in Mice

All animal experiments were performed under protocols approved by the Institutional Animal Care and Use Committee (IACUC) of Daoke Pharmaceutical Technology (Beijing) Co., Ltd. (approval no. IACUC-DKBJ-2024-11-08-02) and conducted in strict compliance with relevant ethical guidelines for animal research.

Round 1: Evaluation of the efficacy of the three prepared mRNA vaccines in mice. Twenty SPF male and twenty SPF female Balb/c mice were randomly divided into four groups, with five mice per group. Mice were vaccinated with 10 μg of GnRH-1 mRNA vaccine, 10 μg of GnRH-2 mRNA vaccine, 10 μg of GnRH-3 mRNA vaccine, or 100 μL of PBS, respectively. A booster immunization was administered 14 days after the primary immunization, followed by a second booster immunization 14 days after the first booster (i.e., on day 28). Serum samples were collected from the orbital venous blood of mice 14 days after the second booster (i.e., on day 42) for serum separation. GnRH-specific antibody levels were determined by indirect ELISA, and sex hormone-related parameters were measured by ELISA. Following euthanasia, gonadal tissues (testes and ovaries) were collected, sectioned, and subjected to histopathological examination.

Round 2: Evaluation of the efficacy of the three prepared mRNA vaccines in mice. Twenty SPF male and twenty SPF female Balb/c mice were randomly divided into four groups, with five mice per group. Mice were vaccinated with 10 μg of GnRH-3 mRNA vaccine, 10 μg of GnRH-4 mRNA vaccine, 10 μg of GnRH-5 mRNA vaccine, or 100 μL of PBS, respectively. A booster immunization was administered 14 days after the primary immunization, followed by a second booster immunization 14 days after the first booster (i.e., on day 28). Serum samples were collected from the orbital venous blood of mice 14 days after the second booster (i.e., on day 42) for serum separation. GnRH-specific antibody levels were determined by indirect ELISA, and sex hormone-related parameters were measured by ELISA. Following euthanasia, gonadal tissues (testes and ovaries) were collected, sectioned, and subjected to histopathological examination.

Round 3: Evaluation of the efficacy of the three prepared mRNA vaccines in mice through mating experiments. Twenty SPF male and sixty SPF female Balb/c mice were randomly divided into four groups, with five males and fifteen females per group. Mice were vaccinated with 10 μg of GnRH-3 mRNA vaccine, 10 μg of GnRH-4 mRNA vaccine, 10 μg of GnRH-5 mRNA vaccine, or 100 μL of PBS, respectively. A booster immunization was administered 14 days after the primary immunization, followed by a second booster immunization 14 days after the first booster (i.e., on day 28). Two days after the second booster (i.e., on day 30), the mice were co-housed, with three female mice and one male mouse per cage. After two days of co-housing, one female mouse was removed from each cage, and a vaginal plug was collected to count the number of fertilized eggs under a microscope. The remaining mice were maintained for further observation to monitor pregnancy and litter size. The experiment was terminated after two months, and mice that had not given birth by that time were considered non-pregnant.

### 2.5 Experimental Design in Cats

Round 1: Dose-dependent effects of GnRH-4 in cats.Twenty male and twenty female cats were randomly divided into four groups, with five cats per group. Cats were vaccinated with 50 μg, 25 μg, or 12.5 μg of GnRH-4 mRNA vaccine, or 1 mL of PBS, respectively. A booster immunization was administered 14 days after the primary immunization. Serum samples were collected from the venous blood of cats 14 days after the booster immunization (i.e., on day 28) for serum separation, and GnRH-specific antibody levels were determined by indirect ELISA. Following euthanasia, gonadal tissues (testes and ovaries) were collected, sectioned, and subjected to histopathological examination.

Round 2: Optimization of immunization schedule in cats.Twenty male and twenty female cats were randomly divided into four groups, with five cats per group. All cats received a primary immunization with 50 μg of GnRH-4 mRNA vaccine. Cats in the single-dose group received no booster immunization. Cats in the two-dose groups received a booster immunization at 14, 21, or 28 days after the primary immunization, respectively. Serum samples were collected from the venous blood of cats 14 days after the booster immunization for serum separation, and GnRH-specific antibody levels were determined by indirect ELISA. Following euthanasia, gonadal tissues (testes and ovaries) were collected, sectioned, and subjected to histopathological examination.

Round 3: Long-term efficacy of GnRH-4 in cats.Thirty male and thirty female cats were randomly divided into six groups, with five cats per group. All cats received a primary immunization with 50 μg of GnRH-4 mRNA vaccine, followed by a booster immunization of the same dose 14 days later. Serum samples were collected from the venous blood of cats at various time points after the primary immunization for serum separation, and GnRH-specific antibody levels were determined by indirect ELISA. Cats were euthanized at 7, 14, 21, 270, and 360 days after the primary immunization, and gonadal tissues (testes and ovaries) were collected, sectioned, and subjected to histopathological examination.

Round 4: Safety evaluation of GnRH-4 in cats.Ten female cats were randomly divided into two groups, with five cats per group. Cats received a single dose of 500 μg of GnRH-4 mRNA vaccine. The general condition and feeding behavior of the cats were observed daily for 14 days. Following euthanasia, organs were collected, photographed, and subjected to histopathological examination.

### 2.6 Western Blot Assay

HEK-293T cells (ATCC CRL-3216) were maintained in Dulbecco’s modified Eagle’s medium (DMEM; Gibco, USA) supplemented with 10% fetal bovine serum (FBS; Excell, Australia) and 1% penicillin-streptomycin(Solarbio, Cat: No.P1400). The cells were incubated under standard culture conditions at 37°C in a humidified atmosphere of 5% COL. For Western blot analysis, HEK-293T cells were transfected with LNPs. After 48 h, cells were collected, and protein concentrations were quantified using a bicinchoninic acid (BCA) assay to ensure equal loading. Subsequently, the protein samples were denatured, separated by SDS-PAGE under reducing conditions, and electrophoretically transferred onto a polyvinylidene difluoride (PVDF) membrane (Thermo Fisher, Cat: 88518). The membrane was then blocked with 5% non-fat milk in PBS containing 0.05% Tween-20 (PBST) to minimize non-specific antibody binding. For immunodetection, the membrane was probed with a mouse monoclonal anti-FLAG primary antibody (TransGen, Cat: HT201-01,1:5000 dilution), followed by extensive washing and incubation with a horseradish peroxidase (HRP)-conjugated goat anti-mouse IgG secondary antibody (TransGen, Cat: HS201-01, 1:10000 dilution). Following additional washes to remove unbound antibodies, specific protein bands were visualized using an enhanced chemiluminescence (ECL) substrate (Beyotime, Cat: P0018S), and the resulting chemiluminescent signals were captured digitally with a Bio-Rad ChemiDoc imaging system.

### 2.7 Detection of serum binding antibodies by ELISA

To quantify antigen-specific antibody responses, an indirect enzyme-linked immunosorbent assay (ELISA) was performed. Briefly, 96-well high-binding microplates were coated with GnRH peptide (EHWSYGLRPG, Tsingke, China) at 100 ng/well in carbonate-bicarbonate coating buffer (pH 9.6) and incubated overnight at 4 °C. Following coating, plates were washed with PBS containing 0.05% Tween-20 (PBST) and blocked with 5% (w/v) non-fat dry milk prepared in PBST for 2 hours at 37 °C to prevent non-specific binding. Serial dilutions of mouse serum samples, prepared in blocking buffer, were added to the plates and incubated for 1 h at 37 °C. After washing, the plates were probed with an HRP-conjugated goat anti-mouse IgG secondary antibody diluted in blocking buffer for 1 h at 37 °C [23]. After a final wash, colorimetric development was initiated with 3,3’,5,5’-tetramethylbenzidine (TMB) substrate (Beyotime, Cat: P0206-100ml), and the enzymatic reaction was terminated with stop solution for TMB substrate (Beyotime, Cat: P0215). Absorbance was immediately measured at 450 nm using a microplate reader, with 630 nm as the reference wavelength (Thermo Fisher, Model: VWD-3400RS). The endpoint antibody titre for each sample was defined as the highest serum dilution factor that yielded an OD450 value exceeding 2.1 times the mean value obtained from negative control sera. The starting dilution was used as the nominal titer for the antibody-negative PBS control group, thus establishing the detection baseline.

### 2.8 Detection of serum parameters

The concentrations of follicle-stimulating hormone (FSH), estradiol (E2), and testosterone (T) in the serum of mice and cats were measured using commercial ELISA kits (all purchased from mlbio, China), and all procedures were performed according to the manufacturer’s instructions.

### 2.9 H&E staining and histopathological scoring

For deparaffinization and rehydration, paraffin sections were sequentially immersed in xylene for 10 min, xylene for another 10 min, anhydrous ethanol for 5 min, 95% ethanol for 5 min, 80% ethanol for 5 min, and 75% ethanol for 5 min, followed by rinsing with tap water. For hematoxylin staining, sections were stained with hematoxylin solution for 5 min, and excess stain was removed by washing with tap water; in cases of overstaining, sections were differentiated for several seconds using differentiation solution, followed by rinsing with tap water for blueing. For eosin staining, sections were stained with eosin solution for 1 min, and excess stain was removed by washing with tap water. For dehydration, clearing, and mounting, sections were sequentially immersed in anhydrous ethanol for 2 min, anhydrous ethanol for another 2 min, anhydrous ethanol for a third 2 min, xylene for 5 min, and xylene for another 5 min. The sections were then removed, an appropriate amount of neutral balsam was applied, and coverslips were placed. Sections were scanned using a slide scanner with a 20× objective lens.

### 2.10 Cryo-TEM Sample Preparation and Imaging

HEK-293T cells were transfected with mRNA-LNPs. After 48Lh, cells were harvested and lysed. The lysate was clarified by centrifugation, and the target proteins were purified using a size-exclusion chromatography column. The purified protein fractions were collected for subsequent cryo-TEM analysis. Cryo-TEM specimen preparation was performed using a Vitrobot Mark IV (Thermo Fisher Scientific). All steps were conducted under strictly controlled environmental conditions: relative humidity was maintained at 100%, and the chamber temperature was held at 10 °C. Quantifoil® R 1.2/1.3, 300-mesh copper grids coated with a continuous carbon film were rendered hydrophilic via glow discharge using a Pelco EasiGlow system (12 mA, 30 s). Immediately after plasma treatment, 3 μL of sample solution was applied onto each grid. Following a 10 s incubation period to ensure uniform spreading and partial dewetting, the grid was blotted for 3 s with Whatman Grade 1 filter paper (Ted Pella) at blotting force setting 1. The grid was then rapidly plunged into liquid ethane cooled by liquid nitrogen to achieve vitrification. Vitrified grids were transferred under liquid nitrogen into cryo-storage boxes and stored in liquid nitrogen until imaging.

Data acquisition was performed on a Thermo Fisher Scientific Tundra cryo-transmission electron microscope operated at 100LkV. Images were recorded at a nominal magnification of ×110,000 with a cumulative electron dose ranges from 20 to 40LeL/Å², using a Thermo Fisher CETA-F direct electron detector.

### 2.11 Transcriptome

Ovaries from female mice and testes from male mice were collected 14 weeks after the completion of the third immunization (day 42) and subjected to transcriptome sequencing, which was performed by BGI. Total RNA was extracted from animal tissues or cultured cells. Tissue samples were homogenized in lysis buffer, whereas cell samples were lysed directly.in the same buffer. Following the addition of chloroform: isopentanol alcohol (24:1) and centrifugation, the supernatant was transferred to a deep-well plate preloaded with binding solution and magnetic beads. Purification was performed using an automated nucleic acid extractor, and the RNA was eluted with RNase-free water. RNA quality was assessed prior to library preparation. For library construction, mRNA was isolated using oligo(dT)-attached magnetic beads, fragmented, and used for first– and second-strand cDNA synthesis. The double-stranded cDNA underwent end repair, A-tailing, adaptor ligation, and PCR amplification. After library quality control, single-stranded circular DNA was generated and sequenced on a DNBSEQ platform by rolling circle amplification and combinatorial Probe-Anchor Synthesis (cPAS). Sequencing data were filtered using SOAPnuke to obtain clean reads. Clean reads were mapped to the reference genome using HISAT2. Gene expression levels were calculated with RSEM. Novel transcripts were predicted using StringTie and Cuffmerge, compared against reference annotation with Cuffcompare, and assessed for coding potential using CPC. Differential gene expression analysis was performed using DESeq2 with genes meeting the criteria of Fold Change ≥ 2 and an adjusted P-value ≤ 0.001 considered significantly differentially expressed. GO enrichment analyses were performed using public databases. Based on the GO annotation results, the differentially expressed genes were functionally categorized. Enrichment analysis for GO terms was performed using the phyper function in R software, with a Q-value ≤ 0.05 considered statistically significant.

### 2.12 Statistical analysis

All experimental data are presented as mean ± standard deviation (SD). Statistical comparisons were performed using GraphPad Prism (v10.0). Significance was determined using one-way analysis of variance (ANOVA) followed by Dunnett’s multiple comparisons test, one-way ANOVA with Tukey’s post hoc test, or two-way ANOVA followed by Fisher’s Least Significant Difference test.as appropriate. A p-value <0.05 was considered statistically significant, with significance levels indicated by *P<0.05, **P<0.01, and ***P<0.001. No data points were excluded from the statistical analysis.

## 3. Results and discussion

### 3.1 Design and in vitro validation of GnRH mRNA vaccines

To obtain GnRH mRNA vaccine candidates with high expression efficiency and strong immunogenicity, this study first conducted a screening of fusion protein scaffolds. Based on three distinct immunoenhancement strategies targeting and prolonged function mediated by the Fc fragment [24, 25], trimeric conformational display mediated by the Foldon domain [26], and particulate self-assembly mediated by the pLS protein [27, 28], three fusion antigens were constructed. Specifically, the GnRH antigen was linked via a flexible linker (GGGGS)L to the Fc fragment (GnRH-1), the Foldon trimerization domain (GnRH-2), and the pLS protein (GnRH-3), respectively (Figure 1A) [29]. The above fusion fragments were cloned into an mRNA vector containing a 5′UTR and a 3′UTR, and the in vitro transcribed mRNAs were encapsulated into lipid nanoparticles (LNPs) composed of SM102, DSPC, cholesterol, and DMG-PEG-2000, yielding three GnRH mRNA-LNP vaccine candidates (Figure 1B). Particle size analysis revealed that the average hydrodynamic diameters of GnRH-1-LNP, GnRH-2-LNP, and GnRH-3-LNP were 101.34 nm, 103.11 nm, and 103.00 nm, respectively, all falling within the optimal size range for efficient LNP delivery (Figure 1C). To validate the expression capabilities of the three LNPs in vitro, each LNP and PBS were incubated with HEK293T cells, and target protein expression was detected by Western blot. The results showed that all three LNPs effectively expressed the corresponding fusion proteins (Figure 1D).

**Figure 1.**
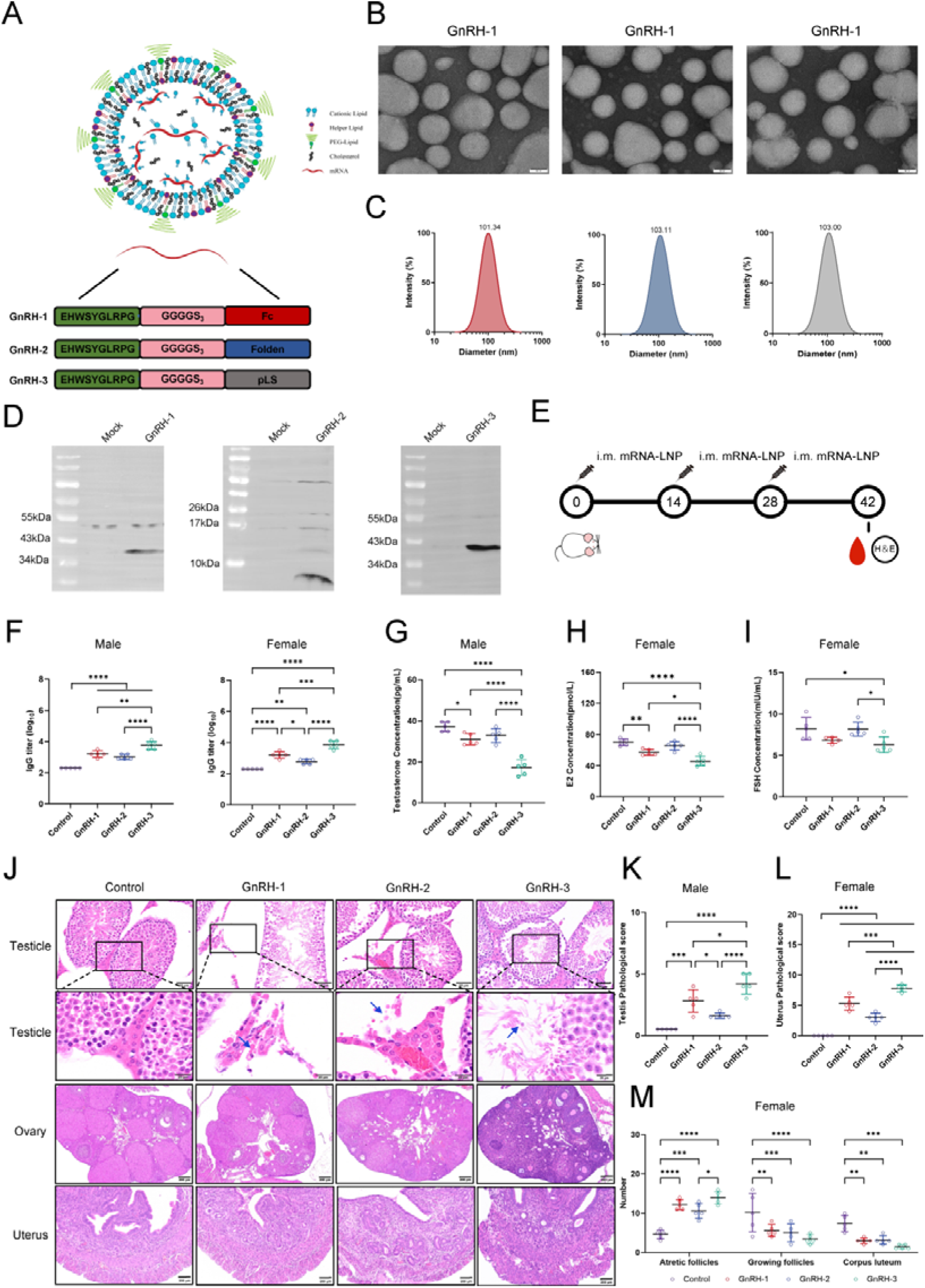
Screening and evaluation of GnRH mRNA vaccine candidates. (A) Schematic of GnRH fusion antigen constructs. GnRH was fused via a flexible linker [(GGGGS)□] to the Fc fragment (GnRH-1), Foldon domain (GnRH-2) or pLS protein (GnRH-3). (B) Transmission electron microscopy image of mRNA. (C) Particle size distributions of GnRH-1-LNP, GnRH-2-LNP and GnRH-3-LNP. (D) Western blot analysis of fusion protein expression in HEK293T cells after incubation with the indicated LNPs. (E) Immunization schedule. (F) GnRH-specific serum IgG titres in female and male mice. (G–I) Serum testosterone (G), estradiol (H) and FSH (I) levels. (J) Representative H&E-stained sections of testis, uterus and ovary. (K,L) Histopathological scores for testis(K) and uterus(L). (M) Quantification of ovarian follicles, including atretic follicles, growing follicles and corpora lutea. Data are presented as mean ± SD. Statistical significance was determined by one-way ANOVA with Tukey’s correction or two-way ANOVA with Fisher’s least significant difference. \**P*<0.05, \*\**P*<0.01, \*\*\**P*<0.001, \*\*\*\**P*<0.0001.

### 3.2 GnRH mRNA vaccines induce durable humoral immunity in mice

It has been reported that serum antibody levels peak at 8–12 weeks after GnRH booster immunization and are significantly higher than those in the control group [30–32]. The above results confirmed that the three GnRH mRNA vaccines effectively expressed the target proteins in vitro. We further evaluated the humoral immune responses induced by these three vaccines in mice and their effects on reproductive organs. To assess the humoral immune responses induced by the vaccines, mice were intramuscularly vaccinated with 10 μg of GnRH-1 mRNA vaccine, GnRH-2 mRNA vaccine, GnRH-3 mRNA vaccine, or PBS (Figure 1E). A booster immunization was administered two weeks after the primary immunization, and serum was collected two weeks after the booster to determine GnRH-specific antibody levels by indirect ELISA. The results showed that compared with the control group, all three vaccine candidate groups generated specific antibodies recognizing the GnRH antigen. Among them, the GnRH-3 immunization group induced significantly elevated antibody levels in both female and male mice compared with the control group (P < 0.0001), and its antibody titers were significantly higher than those in the GnRH-1 (P < 0.01) and GnRH-2 (P < 0.0001) immunization groups (Figure 1F), indicating that the GnRH-3 vaccine exhibited stronger immunogenicity. The effective induction of antibody levels inevitably influences downstream hormone regulation. Hormone level measurements showed that compared with the control group, the GnRH-3 immunization group exhibited significantly decreased concentrations of testosterone, estradiol, and follicle-stimulating hormone (P < 0.05), with greater reductions than those observed in the GnRH-1 and GnRH-2 immunization groups (Figure 1G–I).

Studies have shown that after GnRH administration, male mice in the treatment group exhibited significantly reduced sperm count and testosterone levels, along with seminiferous tubule atrophy; female mice also showed significantly decreased numbers of corpora lutea and reduced levels of sex hormones (progesterone, follicle-stimulating hormone, and luteinizing hormone) [30, 31, 33, 34].Significant alterations in hormone levels ultimately manifest as morphological changes in reproductive organs. To evaluate the effects of the vaccines on reproductive organs, gonadal tissues were collected two weeks after the booster immunization for H&E staining analysis. The results showed that the testicular structure of the control group was normal, with tightly packed and intact seminiferous tubules, well-developed spermatogenic cells at all stages, and abundant sperm within the lumen. In the GnRH-1 and GnRH-2 immunization groups, testicular structure remained largely intact, although some mice exhibited local disruption of seminiferous tubule architecture, a slight reduction in spermatogenic cells, and decreased luminal sperm. The GnRH-3 immunization group displayed more pronounced pathological changes: although testicular structure remained intact, most seminiferous tubules showed structural disorganization, with degeneration and necrosis of spermatogenic cells, a significant reduction in cell numbers, thinning of the tubular wall, empty lumina, and markedly reduced or absent sperm (Figure 1J). Testicular pathology scores were significantly higher in the GnRH-1 and GnRH-3 immunization groups compared with the control group (P < 0.001 and P < 0.0001, respectively); the GnRH-2 group showed an increasing trend. Moreover, the GnRH-3 group exhibited significantly higher scores than the GnRH-1 group (P < 0.05) and the GnRH-2 group (P < 0.0001) (Figure 1K). In female mice, the uteri of mice immunized with all three vaccines exhibited varying degrees of endometrial atrophy and edema of the lamina propria (Figure 1J). Uterine pathology scores were significantly elevated in all three immunization groups compared with the control group (P < 0.0001); additionally, the GnRH-3 immunization group had significantly higher uterine pathology scores than the GnRH-1 group (P < 0.001) and the GnRH-2 group (P < 0.0001) (Figure 1L). Ovarian histological analysis revealed that ovarian structure in the control group was intact, with various stages of growing and mature follicles visible. Compared with the control group, the GnRH-1 and GnRH-2 immunization groups maintained intact ovarian structure but exhibited significantly increased numbers of atretic follicles (P < 0.001 for both) and significantly decreased numbers of growing follicles (P < 0.01 and P < 0.001, respectively) and corpora lutea (P < 0.01 for both). In the GnRH-3 immunization group, although ovarian structure remained intact, the number of atretic follicles was significantly increased (P < 0.0001), while the numbers of growing follicles and corpora lutea were significantly decreased (P < 0.0001 and P < 0.001, respectively) compared with the control group; moreover, the number of atretic follicles was significantly higher than that in the GnRH-2 group (P < 0.05) (Figure 1M). Collectively, based on immunogenicity, hormone levels, and histopathological changes, the GnRH-3 mRNA vaccine outperformed GnRH-1 and GnRH-2 in inducing immune responses and reproductive suppression; therefore, GnRH-3 was selected for further investigation.

### 3.3 Optimization and in vitro validation of GnRH mRNA vaccines

After identifying GnRH-3 as the optimal fusion scaffold, we further optimized the antigen expression module. Previous studies have demonstrated that increasing the copy number of tandem epitopes can effectively enhance vaccine immunogenicity and castration effects [20, 35, 36]. Inspired by this strategy, we performed codon optimization and screened different tandem copy numbers of the GnRH epitope sequence. Specifically, the GnRH epitope sequence was linked in tandem five or ten times via a flexible (GGGGS)L linker and fused to the N-terminus of the pLS protein. Following codon optimization, two candidate antigen modules were obtained, designated GnRH-4 (five tandem repeats) and GnRH-5 (ten tandem repeats), with structural schematics shown in Figure 2A. The above coding sequences were separately cloned into mRNA vectors containing 5′UTR and 3′UTR. After in vitro transcription, the mRNAs were encapsulated into lipid nanoparticles composed of SM102, DSPC, cholesterol, and DMG-2000 to generate GnRH-4-LNP and GnRH-5-LNP. Transmission electron microscopy revealed that both LNPs exhibited uniform spherical morphology, good dispersion, and no obvious aggregation (Figure 2B-C). Particle size analysis showed that the average diameters of GnRH-4-LNP and GnRH-5-LNP were 98.26 nm and 103.00 nm, respectively, both falling within the optimal size range for efficient LNP delivery (Figure 2B-C). To evaluate their expression performance, the two LNPs were transfected into HEK293T cells, and Western blot analysis demonstrated that both effectively expressed the target proteins, with single distinct bands (Figure 2B-C).

**Figure 2.**
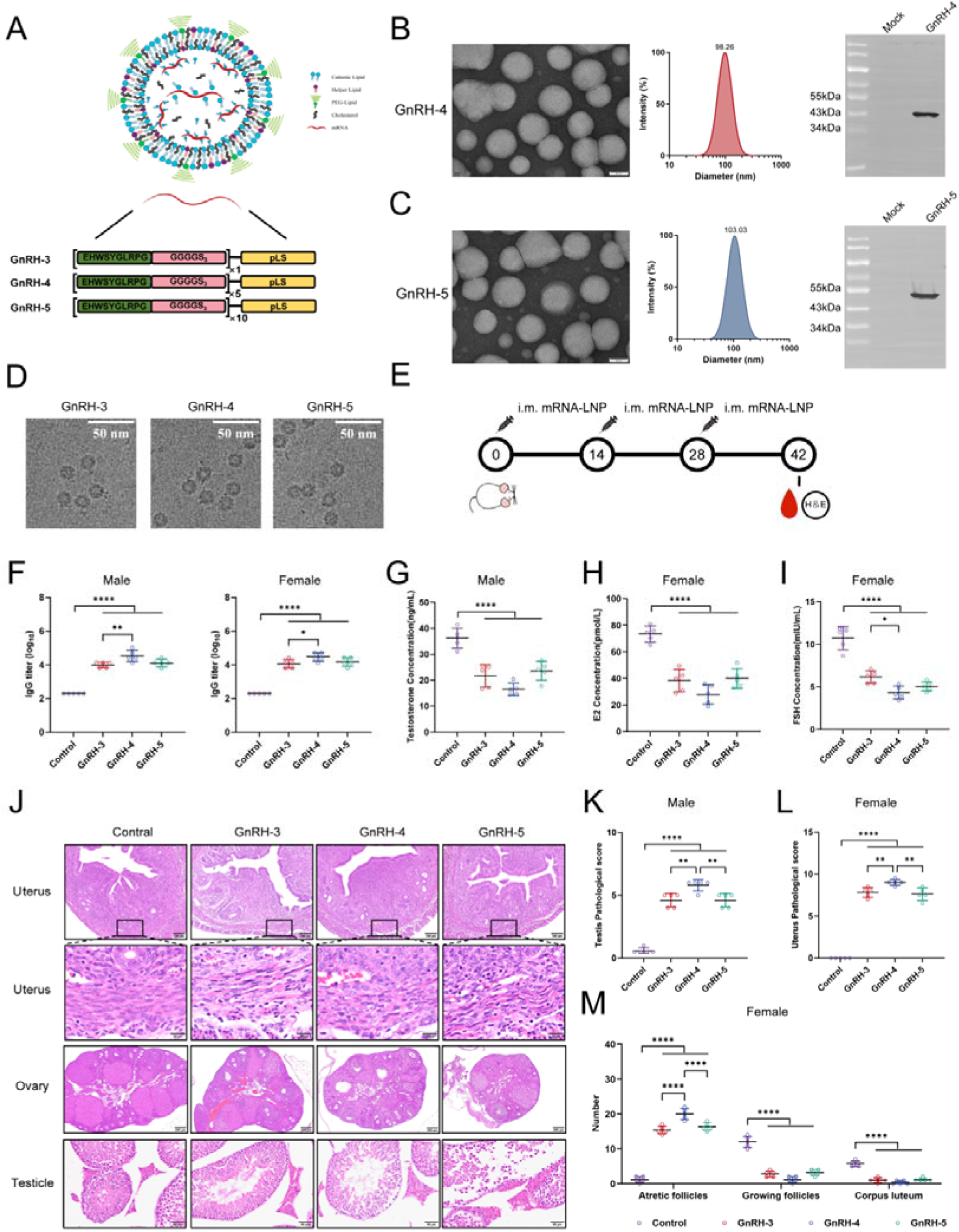
Optimization of tandem GnRH epitopes and evaluation of GnRH-4 and GnRH-5 mRNA-LNP vaccines. (A) Schematic of the GnRH-4 and GnRH-5 antigen constructs, in which five or ten tandem GnRH epitopes were fused to the N terminus of pLS via a flexible linker [(GGGGS)□]. (B,C) Characterization of GnRH-4-LNP(B) and GnRH-5-LNP(C) by transmission electron microscopy and dynamic light scattering. Western blot analysis of antigen expression in HEK293T cells after transfection with the indicated LNPs. (D) Cryo-electron microscopy images of proteins expressed by GnRH-3-LNP, GnRH-4-LNP and GnRH-5-LNP. (E) Immunization schedule. (F) GnRH-specific serum IgG titres in male and female mice. (G–I) Serum testosterone (G), estradiol (H) and follicle-stimulating hormone (FSH; I) levels. (J) Representative H&E-stained sections of testis, uterus and ovary from each group. (K,L) Histopathological scores for testis(K) and uterus(L). (M) Quantification of ovarian follicles, including atretic follicles, growing follicles and corpora lutea. Data are presented as mean ± SD. Statistical significance was determined by one-way ANOVA with Tukey’s correction or two-way ANOVA with Fisher’s least significant difference. \**P*<0.05, \*\**P*<0.01, \*\*\**P*<0.001, \*\*\*\**P*<0.0001.

Furthermore, it has been reported that pLS can form virus-like particle (VLP) structures [18]. To investigate whether the proteins expressed by our LNPs could similarly assemble into VLPs, we employed cryo-electron microscopy to analyze the structural characteristics of the proteins expressed intracellularly by GnRH-1, GnRH-2, and GnRH-3 LNPs. The three LNPs were separately transfected into 293T cells. After 72 h, cells were harvested, and the target protein samples were purified for structural observation by cryo-EM. The results showed that the proteins expressed by all three LNPs formed highly uniform spherical structures in the vitreous ice layer, with diameters concentrated around 10 nm and extremely homogeneous size distribution. No obvious aggregation, fusion, or vesicle formation was observed. A characteristic electron-dense core was visible within the particles, indicating a solid structure, which confirmed that the GnRH-pLS mRNA molecules were stably encapsulated within the cationic lipid matrix rather than forming hollow or multilamellar structures (Figure 2D).

### 3.4 GnRH mRNA-LNPs Induce Durable Humoral Immunity in Mice

Previous studies have demonstrated that the choice of GnRH antigen construct significantly influences the magnitude of the antibody response [37]. To evaluate the immunogenicity of three GnRH mRNA vaccines and their effects on reproductive organs, 40 SPF-grade Balb/c mice were randomly divided into eight groups (n = 5 per group, with equal numbers of males and females). Mice were intramuscularly injected with 10 μg of GnRH-3-LNP, GnRH-4-LNP, GnRH-5-LNP, or empty LNP (control group). The immunization regimen consisted of a primary immunization followed by a booster two weeks later, with a second booster administered after an additional two-week interval. Serum was collected two weeks after the final immunization (Figure 2E), and GnRH-specific antibody levels were measured by indirect ELISA. The results showed that all three vaccine groups generated detectable GnRH-specific antibodies, with levels significantly higher than those in the control group (P < 0.0001). Notably, the GnRH-4 group induced higher antibody titers than both the GnRH-3 and GnRH-5 groups in both male and female mice (males, GnRH-4 vs GnRH-3: P < 0.01; females, GnRH-4 vs GnRH-3: P < 0.05), indicating superior immunogenicity for GnRH-4 (Figure 2F).

The establishment of the aforementioned antibody response further influences endocrine homeostasis, leading to alterations in reproductive hormone levels. Previous studies have confirmed that GnRH vaccines significantly reduce serum testosterone levels [38]. Hormone measurements in this study revealed that serum concentrations of testosterone, estradiol, and follicle-stimulating hormone were significantly decreased in all three vaccine groups compared with the control group (P < 0.0001), with the most pronounced reductions observed in the GnRH-4 group (Figure 2G–I).

Changes in reproductive hormone levels, in turn, induce corresponding morphological alterations in gonadal tissues. It has been previously reported that immunization with a GnRH conjugate significantly arrests spermatogenesis, leading to reduced spermatogenic cell numbers and seminiferous tubule atrophy [31, 39]. To assess the effects of the vaccines on reproductive organs, gonadal tissues were collected two weeks after the final immunization for H&E staining analysis. Testicular histopathology showed that the control group exhibited normal testicular architecture, with abundant and densely packed spermatogenic cells and seminiferous tubules filled with sperm. In the GnRH-3 and GnRH-5 groups, spermatogenic cells were mildly reduced in number and density, and luminal sperm were decreased without complete filling. The GnRH-4 group displayed more pronounced pathological changes, characterized by a significant reduction in spermatogenic cell numbers, thinning of the seminiferous epithelium, markedly decreased cellular density, and a notable decline in luminal sperm (Figure 2J). Testicular pathology scores were significantly higher in all three vaccine groups compared with the control group (P < 0.0001), and the GnRH-4 group exhibited significantly higher scores than both the GnRH-3 and GnRH-5 groups (P < 0.05) (Figure 2K).

Ovarian histological analysis revealed intact ovarian structure in the control group, with abundant growing and mature follicles and no apparent atretic follicles. In all three vaccine groups, the number of atretic follicles was significantly increased (P < 0.0001 vs control), with the GnRH-4 group showing the highest count, which was significantly greater than those in the other two vaccine groups (P < 0.0001). Concurrently, the numbers of growing follicles and corpora lutea were significantly reduced (P < 0.0001 vs control) (Figure 2J, M). Uterine tissues also exhibited varying degrees of atrophy and eosinophilic infiltration, with the most pronounced changes observed in the GnRH-4 group (Figure 2J). Uterine pathology scores were significantly higher in all three vaccine groups compared with the control group (P < 0.0001), and the GnRH-4 group scored significantly higher than the GnRH-3 and GnRH-5 groups (P < 0.01) (Figure 2L).

Collectively, based on immunogenicity, hormone levels, and gonadal histopathological changes, the GnRH-4 mRNA vaccine outperformed GnRH-3 and GnRH-5 in inducing immune responses and reproductive suppression; therefore, GnRH-4 was selected as the lead candidate for subsequent studies.

### 3.5 Evaluation of reproductive suppression efficacy and candidate vaccine screening using a mouse mating model

Pinkham et al. demonstrated that suppression of estrus effectively reduces both the litter rate and litter size in mice. Administration of GnRH to female mice significantly impaired fertility, with only one out of ten females giving birth by day 45 post-treatment [40].To further evaluate the actual impact of the three GnRH mRNA vaccines on reproductive capacity, a mouse mating experiment was designed and conducted. Mice were immunized with 10 μg of GnRH-3 mRNA vaccine, GnRH-4 mRNA vaccine, GnRH-5 mRNA vaccine, or 100 μL of PBS as a control. Male and female mice were co-housed at a ratio of 3:1 for two days, after which one female mouse was randomly selected from each cage for counting of fertilized eggs in the vaginal plug. The remaining female mice were maintained for observation of pregnancy and litter size; mice that had not given birth by the end of the experiment (two months after co-housing) were considered non-pregnant (Figure 3A). The results showed that the number of fertilized eggs in the vaginal plugs of control females was significantly higher than that in the three vaccine groups (P < 0.0001). Notably, the GnRH-4 immunization group exhibited a significantly lower number of fertilized eggs than the GnRH-3 group (P < 0.0001) and the GnRH-5 group (P < 0.05). These findings indicated that the GnRH-4 mRNA vaccine exhibited the most potent suppressive effect on reproductive capacity in mice and could be selected as the preferred candidate for subsequent studies in felines (Figure 3B, C).

**Figure 3.**
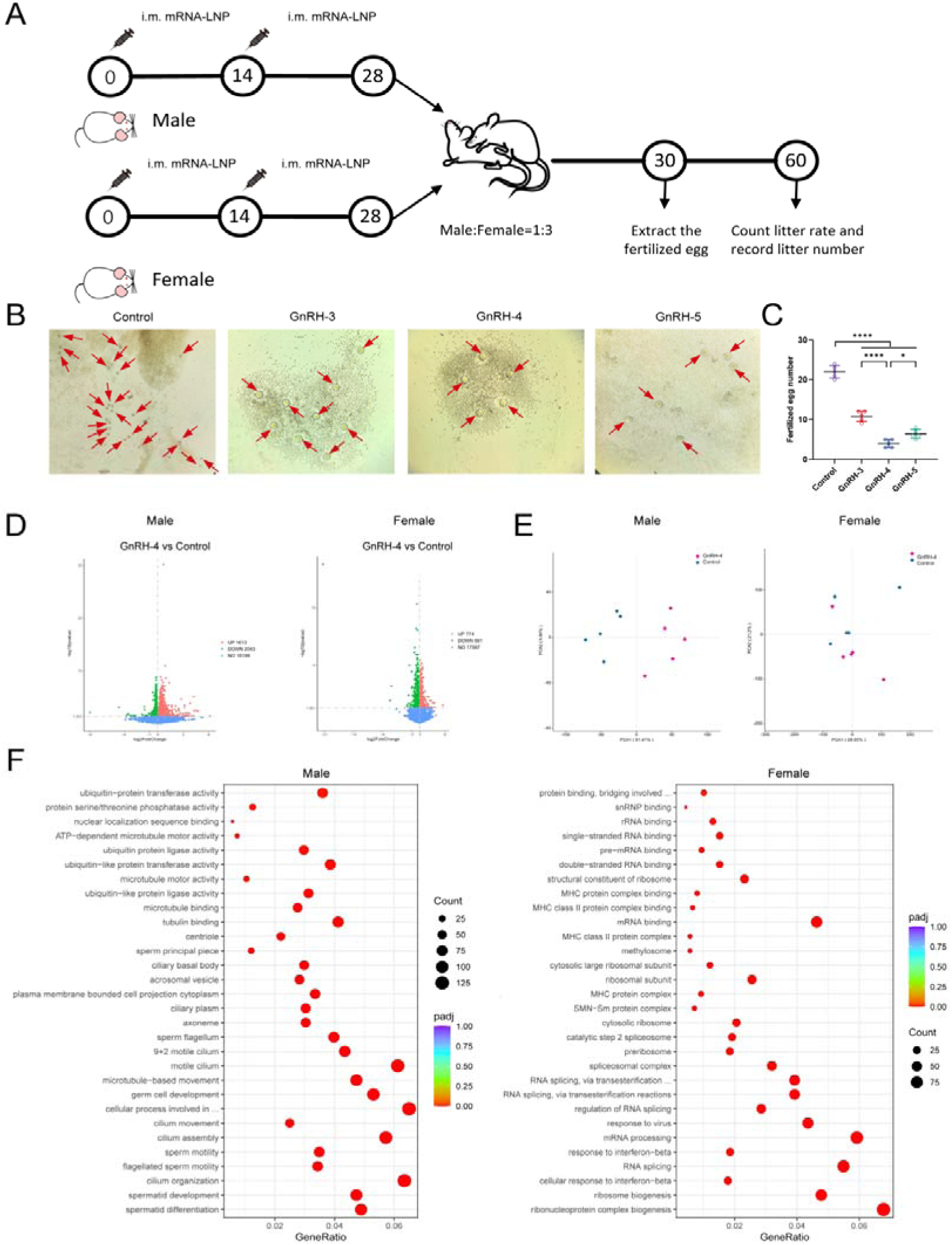
GnRH mRNA vaccines suppress fertility and reshape transcriptional profiles in mice. (A) Schematic of the mating experiment. (B,C) Representative images and quantification of fertilized eggs from control and GnRH vaccine groups. (D) Volcano plots showing differentially expressed genes (DEGs) in GnRH-4-immunized male and female mice. (E) Principal component analysis (PCA) of transcriptomic profiles from GnRH-4-immunized mice. (F) Gene Ontology (GO) enrichment analysis of DEGs in male and female mice. Data are presented as mean ± SD. Statistical significance was determined by one-way ANOVA with Tukey’s correction. \**P*<0.05, \*\**P*<0.01, \*\*\**P*<0.001, \*\*\*\**P*<0.0001.

Litter sizes were statistically analyzed for the control group and the different GnRH treatment groups (Table 1). The results showed that the average litter size in the control group was 8.10 ± 3.75. Compared with the control group, all GnRH treatment groups exhibited a significant reduction in litter size. The GnRH-4 treatment group showed the most pronounced inhibitory effect, with an average litter size of only 0.50 ± 1.08, representing a 93.8% reduction relative to the control group. The GnRH-3 treatment group had an average litter size of 1.60 ± 2.55 (80.2% reduction), and the GnRH-5 treatment group had an average litter size of 2.90 ± 3.18 (64.2% reduction).

**Table 1.** Statistical table of litter size in mice.

| Group | 1 | 2 | 3 | 4 | 5 | 6 | 7 | 8 | 9 | 10 |
| --- | --- | --- | --- | --- | --- | --- | --- | --- | --- | --- |
| Control | 10 | 12 | 8 | 8 | 9 | 7 | 6 | 8 | 13 | / |
| GnRH-3 | 5 | 6 | 5 | / | / | / | / | / | / | / |
| GnRH-4 | 3 | 2 | / | / | / | / | / | / | / | / |
| GnRH-5 | 5 | 7 | 5 | 8 | 4 | / | / | / | / | / |

These results indicated that exogenous GnRH administration significantly inhibited reproductive capacity in mice, with GnRH-4 exhibiting the strongest antifertility effect, followed by GnRH-3, and GnRH-5 showing relatively weaker efficacy. These findings were consistent with the histopathological, antibody, and hormone level results described above.

### 3.6 Transcriptome analysis

Differentially expressed genes (DEGs) were identified by comparing GnRH-4-immunized mice with control mice. In males, 3,656 DEGs were identified, of which 1,613 were upregulated and 2,043 were downregulated. In females, 1,465 DEGs were identified, with 774 upregulated and 691 downregulated (Figure 3D). These DEGs represent key factors involved in GnRH-mediated reproductive suppression in mice. Principal component analysis (PCA) revealed that in male mice, the GnRH-4 group and control group were clearly separated along the PC1 axis, indicating significant differences in the intensity of immune responses between the two groups. In female mice, the separation between the two groups was also notable, although the sample points were more dispersed, suggesting greater inter-individual variability within the group (Figure 3E).

To further infer the main biological functions in immunized mice, GO enrichment analysis was performed on the identified DEGs using the phyper function in R software. As shown in Figure 3F, in male mice, the molecular function category was primarily enriched in ubiquitin-related enzyme activity and microtubule motor activity; the cellular component category was highly enriched in cilium/flagellum-related structures and sperm-specific structures (e.g., sperm principal piece); and the biological process category was significantly enriched in spermatogenesis, motility, and cilium-related processes. In female mice, the molecular function category was mainly enriched in RNA binding and protein binding activities; the cellular component category showed significant enrichment in ribonucleoprotein complexes and spliceosome-related structures; and the biological process category was highly enriched in RNA splicing, ribosome biogenesis, and immune responses. Collectively, these results indicated that in male mice, the effects were primarily directed toward the regulation of germ cell function and motility, whereas in female mice, the effects were more associated with post-transcriptional regulation and immune function.

### 3.7 Immunogenicity and reproductive suppression efficacy of different doses of GnRH-4 mRNA vaccine in cats

Immunization against GnRH has been shown to achieve long-term contraceptive effects in both male and female cats, indicating that the cat is an important model for evaluating GnRH-based interventions [11, 41–44]. To determine the optimal dose of the GnRH-4 mRNA vaccine in the target species (cats), a dose-escalation study was conducted.

Cats were immunized with 50 μg, 25 μg, or 12.5 μg of GnRH-4 mRNA vaccine, or 1 mL of PBS as a control. Serum samples were collected from venous blood 21 days after the booster immunization, and GnRH-specific antibody levels were measured by indirect ELISA (Figure 4A). The results showed that GnRH-specific antibody titers in all three dose groups were significantly higher than those in the control group (P < 0.0001). In both males and females, the 50 μg dose group induced the highest antibody titers. In male cats, the 50 μg group exhibited significantly higher antibody levels than the 12.5 μg group (P < 0.01); in female cats, the 50 μg group also showed significantly higher levels than the 25 μg group (P < 0.05) and the 12.5 μg group (P < 0.01) (Figure 4B).

**Figure 4.**
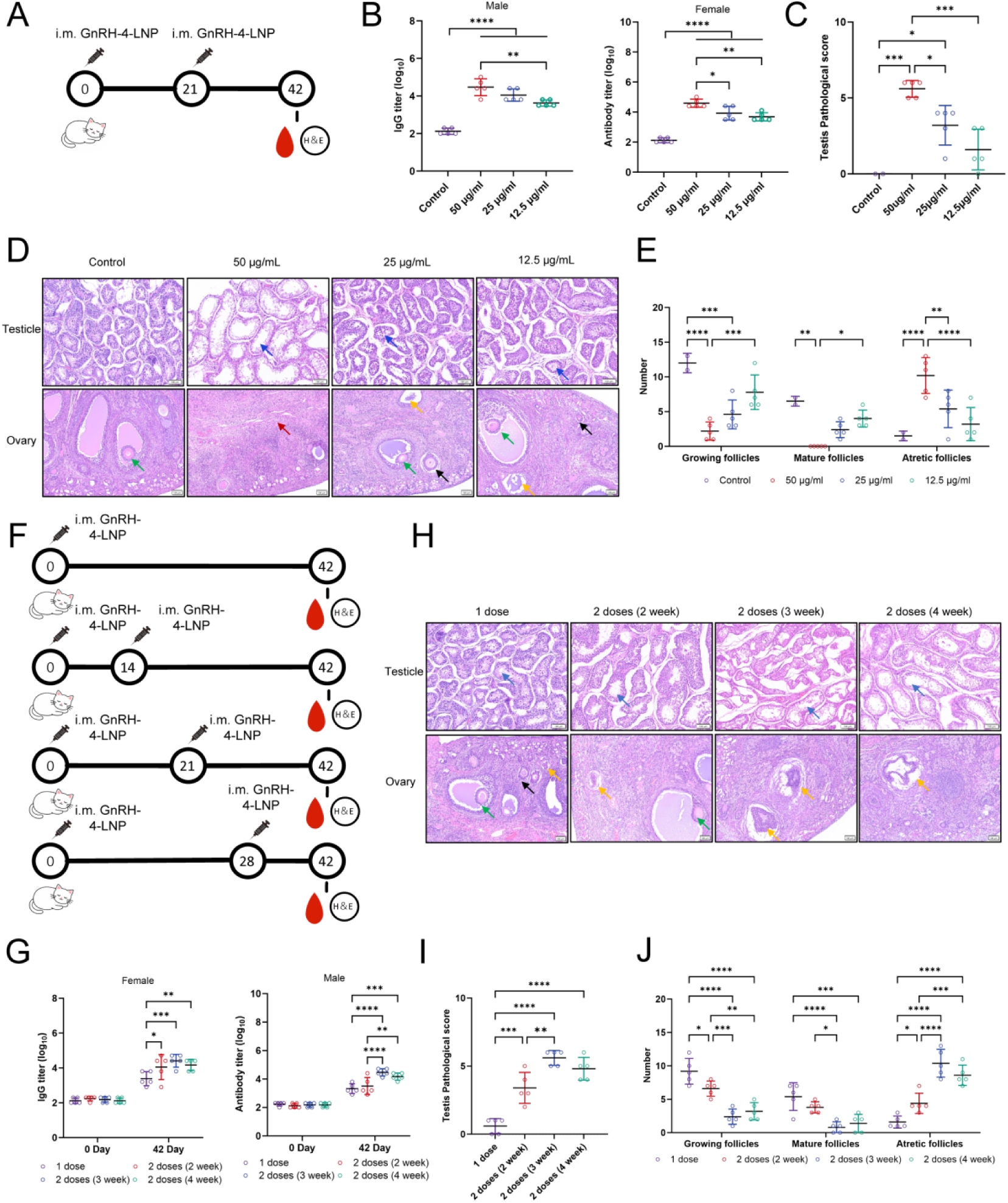
Optimization of GnRH-4 mRNA vaccine dose and immunization schedule in cats. (A) Experimental design for dose comparison. (B) GnRH-specific antibody titres in male and female cats after dose comparison. (C) Semi-quantitative testicular pathology scores after dose comparison. (D) Representative H&E-stained sections of testis and ovary from each dose group. (E) Quantification of ovarian follicles, including growing, mature and atretic follicles. (F) Experimental design for schedule optimization. (G) GnRH-specific antibody titres after schedule optimization. (H) Representative H&E-stained sections of testis and ovary from each immunization schedule. (I) Semi-quantitative testicular pathology scores after schedule optimization. (J) Quantification of ovarian follicles after schedule optimization. Data are presented as mean ± SD. Statistical significance was determined by one-way ANOVA with Tukey’s correction or two-way ANOVA with Fisher’s least significant difference. \**P*<0.05, \*\**P*<0.01, \*\*\**P*<0.001, \*\*\*\**P*<0.0001.

To evaluate the effects of the vaccine on reproductive organs, histopathological examination of gonadal tissues was performed. Animals were euthanized following immunization, and testicular and ovarian tissues were collected for histopathological examination (Figure 4A). In male cats, semi-quantitative testicular pathology scoring revealed that the 50 μg group achieved the highest scores, with extremely significant differences compared with the control group and the 12.5 μg group (both P < 0.001), and a significant difference compared with the 25 μg group (P < 0.05). The 25 μg group also exhibited significantly higher scores than the control group (P < 0.05) (Figure 4C). All immunization groups displayed varying degrees of testicular damage, characterized by reduced seminiferous tubule diameter, thinning of the seminiferous epithelium, and decreased sperm numbers, with the most severe lesions observed in the 50 μg group, followed by the 25 μg group, and the mildest changes in the 12.5 μg group (Figure 4D). In female cats, compared with the control group, all immunization groups exhibited physiological changes in ovarian tissue: the number of growing follicles was significantly reduced, with the 50 μg and 25 μg groups showing significant decreases compared with the control group (P < 0.0001 and P < 0.001, respectively), and the 12.5 μg group also showing a decreasing trend; moreover, the 50 μg group exhibited a significantly greater reduction than the 12.5 μg group (P < 0.001). The number of mature follicles also showed a decreasing trend, with the 50 μg group demonstrating significant reductions compared with both the control group and the 12.5 μg group (P < 0.01 and P < 0.05, respectively). The number of atretic follicles was significantly increased, with all three dose groups showing an increasing trend compared with the control group; notably, the 50 μg group exhibited an extremely significant increase compared with the control group and the 12.5 μg group (P < 0.0001), and a significant increase compared with the 25 μg group (P < 0.01) (Figure 4D, E).

Based on the combined assessment of antibody responses and reproductive organ pathological changes, the 50 μg dose exhibited the most pronounced effects in terms of inducing immune responses and suppressing reproductive function and was therefore selected as the optimal dose for subsequent experiments.

### 3.8 Effects of Different Booster Immunization Time Points on the Efficacy of GnRH-4 mRNA Vaccine

Multiple studies have demonstrated that the interval between vaccine doses significantly influences the immune response. Clinical trials of the ChAdOx1 nCoV-19 (AZD1222) vaccine showed that extending the interval between the two doses to 12 weeks or longer increased vaccine efficacy from 55.1% to 81.3%, and antibody levels were more than doubled. In another study, extending the dosing interval of the BNT162b2 mRNA vaccine to 6–14 weeks, compared with a 3– to 4-week schedule, significantly enhanced neutralizing antibody titers. These findings suggest that a longer interval can enhance immune responses and that appropriate adjustment of the dose interval can optimize individual immune outcomes [45–47]. Having established 50 μg as the optimal dose, this study further optimized the immunization schedule, focusing on the impact of different booster time points. The single-dose group received no booster immunization, while the two-dose groups received a booster immunization on day 14, day 21, or day 28 after the primary immunization (Figure 4F).

To evaluate the humoral immune responses induced by different immunization regimens, serum was collected after the booster immunization, and GnRH-specific antibody levels were measured by indirect ELISA. Antibody detection results showed that all booster groups exhibited significantly higher antibody titers than the single-dose group. Among them, the day 21 booster group induced the highest antibody levels in both females (P < 0.001) and males (P < 0.0001), followed by the day 28 booster group (females: P < 0.01; males: P < 0.001), while the day 14 booster group showed relatively lower levels (females: P < 0.05). Notably, in male cats, the day 21 booster group exhibited significantly higher antibody levels than the day 14 booster group (P < 0.0001) (Figure 4G).

Histopathological evaluation was subsequently performed to further assess the impact of different booster schedules on reproductive organ morphology. Animals were euthanized following immunization, and testicular and ovarian tissues were collected for histopathological examination. In male cats, the day 21 booster group exhibited the most severe testicular damage, characterized by reduced seminiferous tubule diameter, significant thinning of the tubular wall, and a marked decrease in spermatogenic cell numbers. The day 28 and day 14 booster groups also displayed similar pathological changes, albeit to a lesser extent (Figure 4H). Testicular pathology scores indicated that all two-dose groups had significantly higher scores than the single-dose group (P < 0.001), and the day 21 booster group had significantly higher scores than the day 14 booster group (P < 0.01) (Figure 4I). In female cats, compared with the single-dose group, all booster groups exhibited significant reductions in the numbers of growing and mature follicles and significant increases in the number of atretic follicles. Regarding growing and mature follicles, the day 21 and day 28 booster groups showed the most significant reductions compared with the single-dose group (P < 0.001), the day 14 booster group also showed a decreasing trend, and the day 21 booster group had significantly lower numbers than the day 14 booster group (P < 0.05). Regarding atretic follicles, the day 21 and day 28 booster groups showed the most significant increases compared with the single-dose group (P < 0.0001), and both groups had significantly higher numbers of atretic follicles than the day 14 booster group (P < 0.001) (Figure 4H, J).

Collectively, based on antibody levels and histopathological parameters, the two-dose immunization regimen was superior to the single-dose regimen; among the two-dose schedules, boosting on day 21 after the primary immunization yielded the best efficacy, followed by boosting on day 28, with boosting on day 14 showing relatively weaker effects.

### 3.9 Sustained Immune Responses and Reproductive Suppression Induced by GnRH-4 mRNA Vaccine in Cats

The long-term durability of a vaccine is a critical indicator for evaluating its protective efficacy. A long-term study on the herpes zoster vaccine demonstrated that its protective efficacy against the disease remained significant for up to 10 years after vaccination [48] To assess the durability of the immune response induced by the GnRH-4 mRNA vaccine in cats, a 12-month long-term efficacy study was conducted. Both male and female cats were immunized according to the optimized regimen (50 μg primary immunization, booster on day 21). Serum samples were collected at various time points after primary immunization to measure antibody levels; animals were euthanized at different time points for histopathological examination of gonadal tissues (Figure 5A).

**Figure 5.**
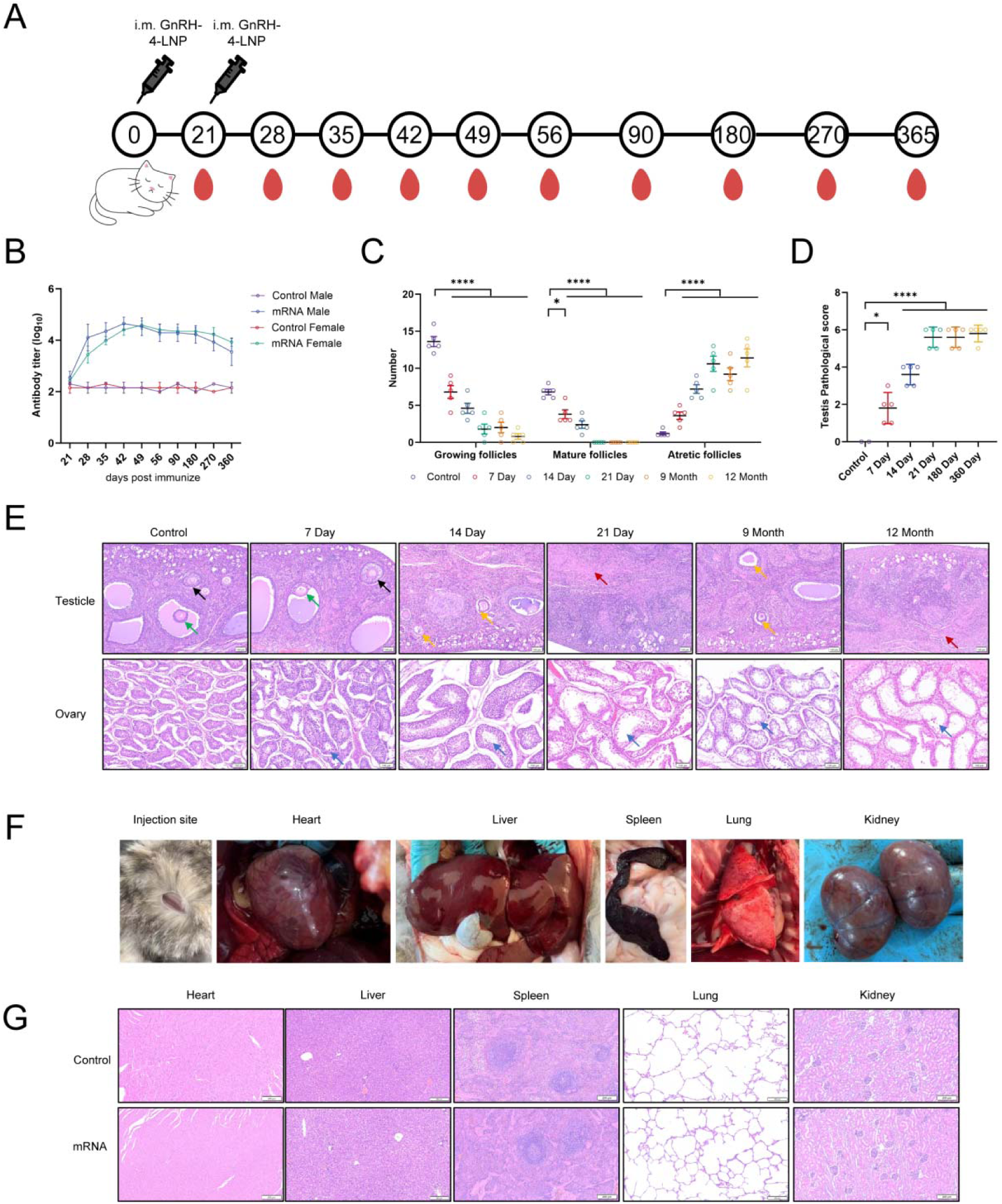
Long-term efficacy and biosafety evaluation of the GnRH-4 mRNA vaccine in cats. (A) Experimental timeline for the 12-month durability study. (B) Kinetics of GnRH-specific antibody titres in male and female cats after immunization. (C) Quantification of ovarian follicles at the indicated time points during long-term follow-up,including growing, mature and atretic follicles. (D) Semi-quantitative testicular pathology scores at the indicated time points. (E) Representative HE-stained sections of testes and ovaries from each group of cats at each time point. (F) Gross examination of the injection site and major organs after high-dose GnRH-4 mRNA vaccine administration. (G) Representative H&E-stained sections of major organs, including heart, liver, spleen, lung and kidney, after high-dose vaccination. Data are presented as mean ± SD. Statistical significance was determined by one-way ANOVA with Tukey’s correction or two-way ANOVA with Fisher’s least significant difference. \**P*<0.05, \*\**P*<0.01, \*\*\**P*<0.001, \*\*\*\**P*<0.0001.

Antibody kinetics revealed that serum GnRH-specific antibody levels in immunized cats were significantly higher than those in the control group starting from day 21 after primary immunization. Antibody titers exhibited a sustained upward trend: in male cats, titers peaked on day 42, then slightly declined but remained stable and consistently elevated compared with the control group; in female cats, titers peaked on day 49 and subsequently maintained a high plateau phase, remaining persistently higher than those in the control group (Figure 5B).

The durability of the immune response ultimately required validation at the histological level. To evaluate the long-term effects of the vaccine on reproductive organs, testicular and ovarian tissues were collected at various time points for H&E staining analysis. In male cats, testicular tissues at all time points exhibited varying degrees of pathological changes compared with the control group. Among these, the most severe testicular damage was observed on day 21, month 9, and month 12, characterized by a marked reduction in spermatogenic cell numbers within the seminiferous tubules; similar but milder changes were observed on day 7 and day 14 (Figure 5E). Testicular pathology scores on day 14, day 21, month 9, and month 12 were extremely significantly higher than those in the control group (P < 0.0001), and the score on day 7 was also significantly higher than that in the control group (P < 0.05) (Figure 5D). In female cats, ovarian tissues on day 7, day 14, day 21, month 9, and month 12 all exhibited consistent physiological changes: compared with the control group, the numbers of growing and mature follicles were extremely significantly reduced (P < 0.0001), and the number of atretic follicles was extremely significantly increased (P < 0.0001) (Figure 5C, E).

Collectively, these results indicate that following a single booster immunization, the GnRH-4 mRNA vaccine induced immune responses and reproductive suppression effects lasting at least 12 months in cats, demonstrating excellent long-term efficacy.

### 3.10 Biosafety assessment of a high dose of GnRH-4 mRNA vaccine in cats

Previous studies have shown that GnRH vaccines can cause adverse effects in animals. It has been reported that elephants developed local swelling, stiffness, and mild lameness after vaccination with a GnRH vaccine, and white-tailed deer also exhibited abscess formation following primary immunization [48].To evaluate the clinical safety of the GnRH-4 mRNA vaccine, a high-dose tolerance study was conducted. Female cats received a single intramuscular injection of 500 μg of GnRH-4 mRNA vaccine. The general condition and feeding behavior of the cats were observed daily for 14 days. Subsequently, cats were euthanized, and major organs were collected for gross observation and histopathological examination.

The results showed that all immunized cats remained in good mental status with normal appetite throughout the observation period, and no abnormal clinical manifestations were observed. Gross examination revealed normal organ morphology with no evidence of congestion, edema, or necrosis (Figure 5F). Histopathological examination showed that the structural integrity of major organs, including the heart, liver, spleen, lungs, and kidneys, was maintained, with normal cellular morphology and no inflammatory infiltration, degeneration, necrosis, or other pathological changes (Figure 5G).

Collectively, these results indicate that even at a dose ten times higher than the optimal immunization dose, the GnRH-4 mRNA vaccine did not induce any observable clinical toxicity or histopathological damage, demonstrating its favorable biosafety profile.

## 4. Conclusion

In this study, we developed a novel GnRH mRNA vaccine using lumazine synthase nanoparticles as a carrier and evaluated its immunogenicity, reproductive suppression efficacy, and safety in mice and cats. Through systematic optimization, GnRH-4, containing five tandem GnRH epitopes fused to pLS, was identified as the optimal candidate. In mice, GnRH-4 induced robust antibody responses, significant gonadal atrophy, and a marked reduction in litter size. Transcriptomic analysis revealed sex-specific mechanisms, with male mice showing enrichment in spermatogenesis-related pathways and females in RNA splicing and immune responses. In cats, the vaccine exhibited dose-dependent immunogenicity, with an optimal regimen of 50 μg administered as a two-dose schedule with a 21-day interval. Notably, GnRH-4 induced sustained immune responses and reproductive suppression lasting at least 12 months, and high-dose administration confirmed excellent biosafety. Collectively, our findings establish the pLS-based GnRH mRNA vaccine as a safe, effective, and long-lasting immunocastration strategy, offering a promising alternative to traditional castration methods for animal management.

## Data Availability

The data that support the findings of this study are available from the corresponding author upon reasonable request.

## Founding

This research did not receive any funding

